# Volumetric grey matter alterations in adolescents and adults born very preterm suggest accelerated brain maturation

**DOI:** 10.1101/127365

**Authors:** Vyacheslav R. Karolis, Sean Froudist-Walsh, Jasmin Kroll, Philip J. Brittain, Chieh-En Jane Tseng, Kie-Woo Nam, Antje A. T. S. Reinders, Robin M. Murray, Steven C. R. Williams, Paul M. Thompson, Chiara Nosarti

**Affiliations:** Department of Psychosis Studies, Institute of Psychiatry, Psychology and Neuroscience, King’s College London, London, UK; Department of Neuroscience, Friedman Brain Institute, Icahn School of Medicine at Mount Sinai, New York, NY, 10029, USA; Centre for Neuroimaging Sciences, Institute of Psychiatry, Psychology and Neuroscience, King’s College London, London, UK; Imaging Genetics Center, Mark and Mary Stevens Institute for Neuroimaging and Informatics, Keck School of Medicine of USC, University of Southern California, Marina del Rey, CA, USA; Centre for the Developing Brain, Division of Imaging Sciences & Biomedical Engineering, King’s College London, London, UK

**Keywords:** brain development, neuroanatomy, plasticity, outcome studies, associational areas

## Abstract

Previous research investigating structural neurodevelopmental alterations in individuals who were born very preterm demonstrated a complex pattern of grey matter changes that defy straightforward summary. Here we addressed this problem by characterising volumetric brain alterations in individuals who were born very preterm from adolescence to adulthood at three hierarchically related levels - global, modular and regional. We demarcated structural components that were either particularly resilient or vulnerable to the impact of very preterm birth. We showed that individuals who were born very preterm had smaller global grey matter volume compared to controls, with subcortical and medial temporal regions being particularly affected. Conversely, frontal and lateral parieto-temporal cortices were relatively resilient to the effects of very preterm birth, possibly indicating compensatory mechanisms. Exploratory analyses supported this hypothesis by showing a stronger association of lateral parieto-temporal volume with IQ in the very preterm group compared to controls. We then related these alterations to brain maturation processes. Very preterm individuals exhibited a higher maturation index compared to controls, indicating accelerated brain ageing and this was specifically associated with younger gestational age. We discuss how the findings of accelerated maturation might be reconciled with evidence of delayed maturation at earlier stages of development.

## Introduction

Numerous studies show widespread structural grey matter alterations associated with very preterm birth (< 32 weeks of gestation) at various developmental stages (Ball et al., 2012; de Kieviet, Zoetebier, Van Elburg, Vermeulen, & Oosterlaan, 2012; Nosarti et al., 2008). We previously suggested that differences in structural measures between very preterm born individuals and controls may decrease over time (Nam et al., 2015; Nosarti et al., 2014), possibly reflecting delayed maturation and subsequent “catch-up” later in development. However, the complex pattern of both increases and decreases in cortical volume and thickness (Bjuland, Lohaugen, Martinussen, & Skranes, 2013; Martinussen et al., 2005; Nosarti et al., 2008; Peterson et al., 2000) defy an easy generalization with respect to their developmental significance. Even in typically developing children, longitudinal changes in cortical volumes (or thickness) are not one-directional (e.g. always decreasing) or uniform across the brain and reach growth peaks at different ages (Gogtay et al., 2004; Sowell et al., 2004). Consequently, the possibility of developmental delay following very preterm birth warrants appropriate quantitative analyses.

Here we will address two gaps in our current understanding of the long-lasting developmental sequelae of very preterm birth. Firstly, we will detail the longitudinal brain trajectories of grey matter volume (GMV) and inter-hemispheric volume lateralisation in very preterm individuals from adolescence (age 15) to adulthood (age 30). This will extend prior findings reported in the same cohort (Nosarti et al., 2002; Nosarti et al., 2008; Nosarti et al., 2014), both in terms of wider age range and in terms of how brain structural characteristics are defined. Secondly, we will relate these alterations to maturational processes and assess whether specific GMV trajectories indicate delayed or accelerated maturation in very preterm individuals compared to controls.

Mass-univariate analytic approaches, such as voxel-based morphometry (Ashburner & Friston, 2000), are invaluable tools to map the spatial extent and location of volumetric differences. However, these approaches neglect the fact that regional morphological features are interdependent, due to common genetic and environmental factors (Evans, 2013). Individuals who experience neurodevelopmental impairments, including those born very preterm, demonstrate altered patterns of structural co-variance (Modinos et al., 2009; Nosarti, Mechelli, et al., 2011; Scheinost, Kwon, et al., 2015). Here, we delineate the morphological profile of the very preterm brain in terms of alterations in structural brain ‘modes’ (Douaud et al., 2014; O’Muircheartaigh et al., 2014), taking into account both regional volumetric measures and their interindividual covariance. We refine this approach by defining structural markers that summarise GMV characteristics at different levels of abstraction - starting from segregated brain regions to brain networks (modules) to whole-brain volume, as shown in Figure 1*A*.

We formulate two contrasting hypotheses. In line with a developmental ‘catch-up’ hypothesis, maturational GMV patterns in very preterm adolescents would be similar to those found in *younger* controls (Franke, Luders, May, Wilke, & Gaser, 2012), but the discrepancy between the two groups would diminish later. Alternatively, maturational GMV patterns in very preterm individuals could be similar to those found in *older* controls. This second hypothesis is based on evidence relating grey matter alterations in the very preterm brain to the underdevelopment of white matter connectivity (Ball et al., 2012; Pandit et al., 2014), possibly due to neonatal brain injury affecting the development of pre-myelinating oligodendrocytes and subplate neurons (Back, Riddle, & McClure, 2007). White matter loss is typical in ageing (Gunning-Dixon, Brickman, Cheng, & Alexopoulos, 2009), therefore the developmental phenotype associated with GMV alterations in very preterm born individuals may mirror patterns that reflect accelerated ageing in white matter.

## Results

### Sample characteristics

Study sample characteristics are shown in Table 1. Very preterm participants were slightly older than controls (F(1,554) = 4.6, p = .032). There was no significant between-group difference in sex (p > .28) and handedness (p > .4). Finally, there was no significant between-group difference in parental socio-economic status at adolescence and young adulthood (both p >.3), although the two groups differed at adulthood (χ^2^ (2) = 9.82, p < .05), with more preterm participants belonging to lower socio-economic groups.

**Table 1.** Sample statistics

|  | <u>ADOLESCENCE</u> |  | <u>EARLY ADULTHOOD</u> |  | <u>ADULTHOOD</u> |  |
| --- | --- | --- | --- | --- | --- | --- |
|  | Control | Preterm | Control | Preterm | Control | Preterm |
| **N** | 92 | 160 | 53 | 72 | 87 | 96 |
| **Age (SD)** | 15.1 (.9) | 15.3 (.5) | 19.5 (1.4) | 20.1 (1.3) | 30.3 (4.6) | 30.8 (2.3) |
| **Male ratio** | 0.57 | 0.51 | 0.49 | 0.49 | 0.46 | 0.58 |
| **Gestation Age (SD)** | NA | 29.1 (2.2) | NA | 28.8 (2.3) | NA | 29.2 (2.2) |
| **Birth Weight (SD)** | NA | 1275 (367) | NA | 1225 (370) | NA | 1296 (371) |
| **Socio-economic status (%)** |  |  |  |  |  |  |
| **I-II** | 42.4 | 41.3 | 41.5 | 43.1 | 65.5 | 53.1 |
| **III** | 32.6 | 40.6 | 26.4 | 44.5 | 18.4 | 38.5 |
| **IV-V** | 18.5 | 16.9 | 13.2 | 9.8 | 2.3 | 8.3 |
| **Unclassified/ Missing** | 6.5 | 1.2 | 18.9 | 2.6 | 13.8 | 0 |
| **Handedness (%)** |  |  |  |  |  |  |
| **Right** | 80.4 | 80.0 | 75.5 | 83.3 | 77.0 | 88.5 |
| **Left** | 8.7 | 15.0 | 5.7 | 16.7 | 6.9 | 10.4 |
| **Ambidextrous** | 0 | 0 | 0 | 0 | 2.3 | 0 |
| **Missing** | 10.9 | 5.0 | 18.9 | 0 | 13.8 | 1.0 |

Socio-economic status categories are in accordance with the Standard Occupational Classification 1980 (SOC1980).

### Module partitioning using normative data set

Hierarchical structural markers, incorporating both raw volumetric measures and their inter-individual covariance, were constructed based on an independent normative data set (**Materials and Methods** and Supplementary Materials), using a combination of modularity analysis and principal components analysis **(Materials and Methods** and Figure 1*B*). Three hierarchical levels were considered: global, modular and regional. Two independent hierarchies were created, one for the means of homologous regions in the left and right hemisphere, GMVs, and the other for the difference between homologous regions in the left and right hemisphere, hereafter - lateralisation indices (LIs). Importantly, the present hierarchical construct endows markers at a lower level with values that are relative with respect to markers at a higher level. Broadly speaking, higher components in the hierarchy (brain ‘modes’) characterise generic motifs persistent across a subset of lower structural components; correspondingly, lower components characterise alterations, which are over and above alterations captured by a higher-level brain ‘mode’. We refer to these markers as module- and region-specific to distinguish them from raw volumetric markers.

**Figure 1.**
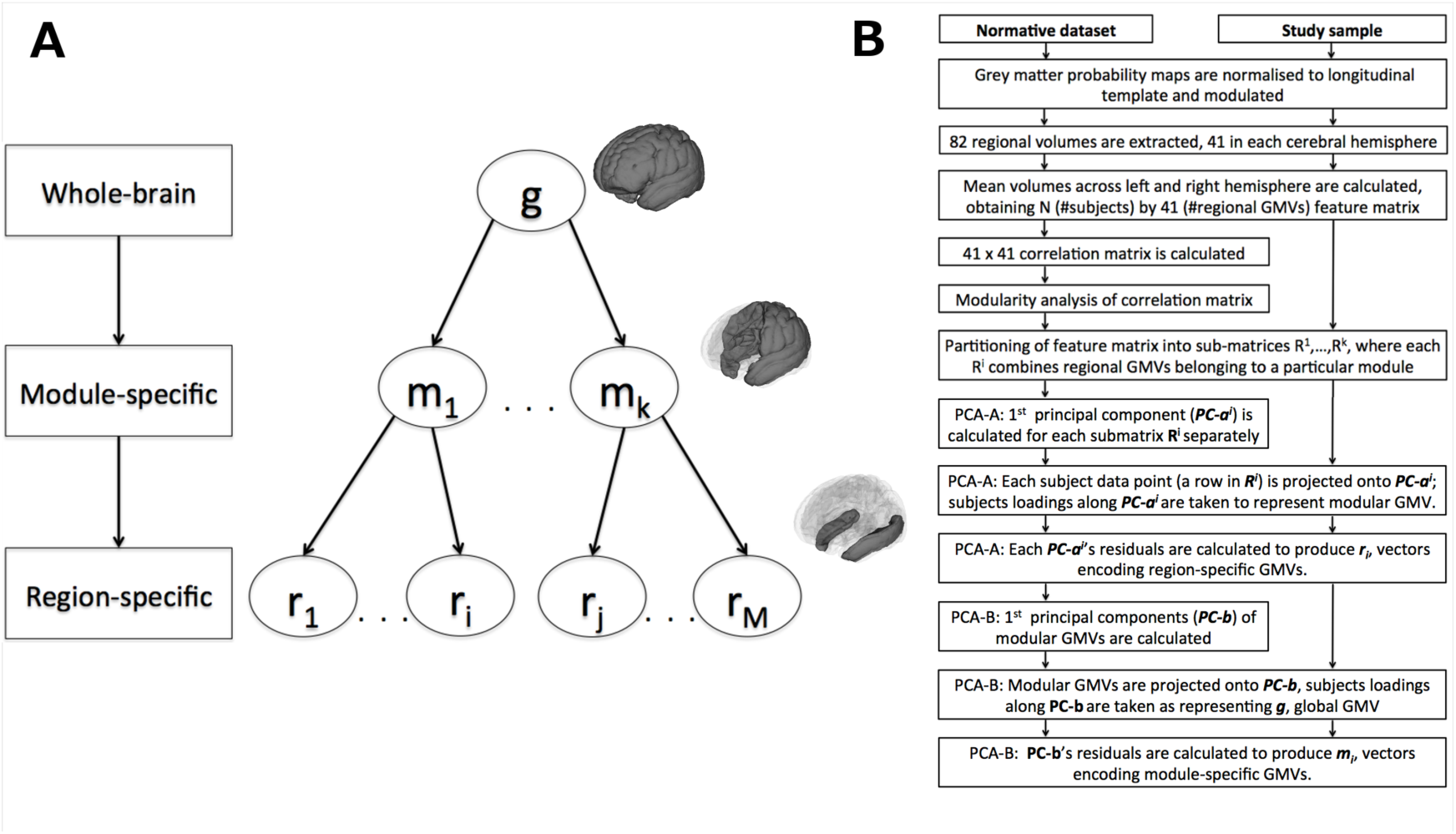
Data structure and feature re-parameterisation. (a) Hierarchy of structural markers. (b)Feature re-parameterisation flow-chart.

Module partitioning for GMVs and LIs is shown in Figure 2. Four modules were obtained for GMVs and 5 modules were obtained for LIs. Structural modules were organised in spatially consistent units. Modular GMVs explained 45 to 60% of variance in regional GMVs in the normative data set, whereas modular LIs explained 24 to 52% of variance. There was a greater similarity between GMV modules than between LI modules, with 70% and 33% of the variance in modular GMVs and LIs explained by global GMV and LI factors, respectively. Given the GMV modules’ very distinctive spatial organisation, we will refer to them, in exact order, as lateral parieto-temporal, posterior medial, subcortical/medial temporal and fronto-striatal modules.

**Figure 2.**
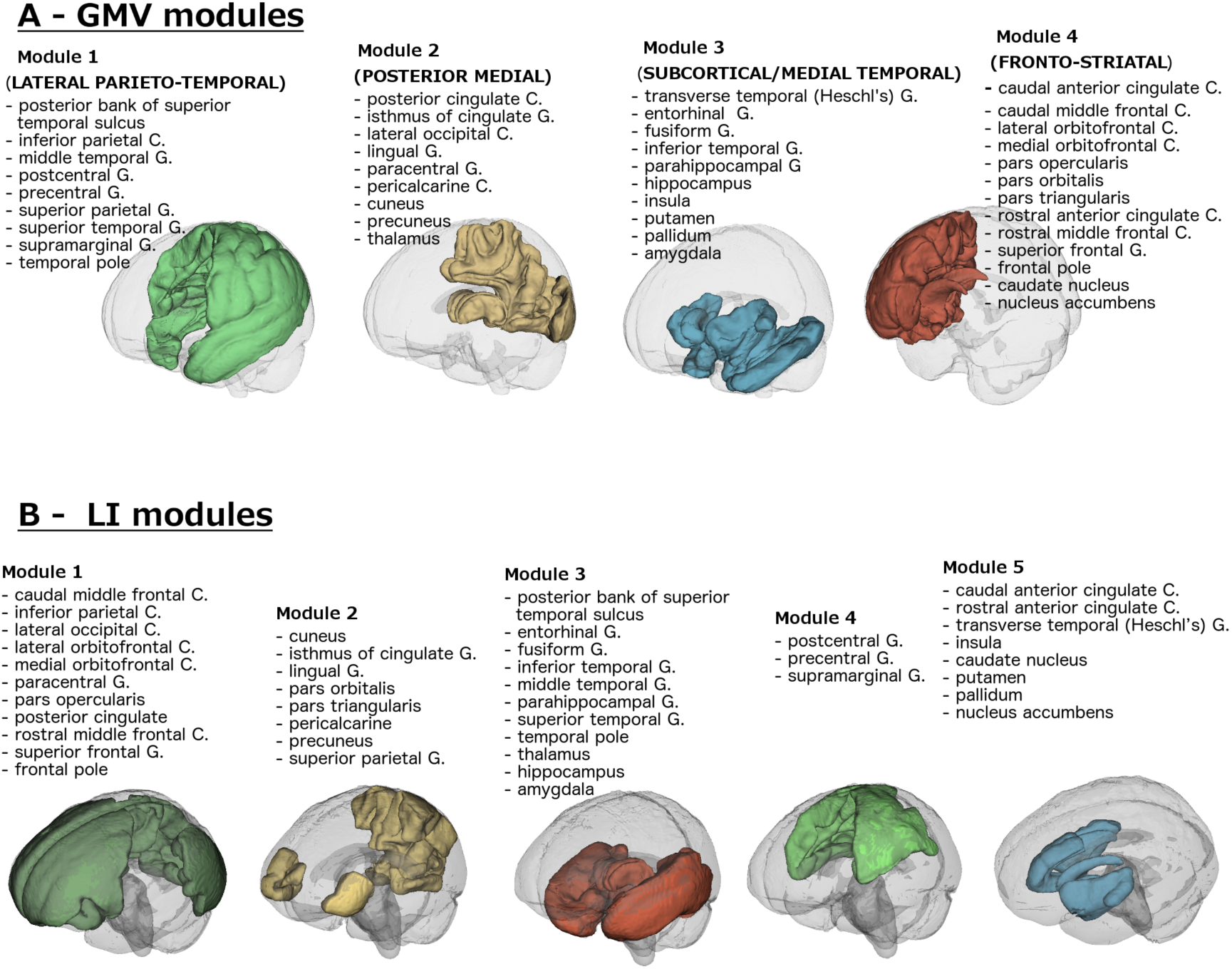
Modular partitioning in the normative data set. (a) On the basis of grey matter volumes, average across left and right. (b) On the basis of lateralisation indices, i.e., the difference in volumes between left and right hemisphere.

## Grey matter volume alterations

The effect of group and its interaction with time of assessment in each GMV on longitudinal GMV trajectories were accounted for by mixed-effect regression models, which also included variables of no interest and intercept and participants’ age as random factors (see **Materials and Methods** for details). Statistics for the main effect of group on GMVs are shown in Table 2.

**Table 2.** Summary of GMV alterations in very preterm born individuals compared to controls.

|  | **Standardised $\beta$ [95% CI]** | **t (df = 552)** |
| --- | --- | --- |
| **Global** | -0.24 [-.42 -.05] | 2.56 |
| **Module-specific** |  |  |
| *Lateral Parieto-Temporal module* | .40 [0.17 0.63] | 3.36 |
| *Subcortical / Medial Temporal module* | -.83 [-1.04 -.62] | 7.76 |
| *Fronto-Striatal module* | .43 [0.25 0.61] | 4.73 |
| **Region-specific** |  |  |
| *Caudate Nucleus* | -.57 [-.80 -.35] | 4.97 |
| *Posterior Superior Temporal Sulcus* | -.47 [-.72 -.21] | 3.60 |
| *Caudal Part of Anterior Cingulate Gyrus* | -.47 [-.73 -.21] | 3.58 |
| *Inferior Parietal Cortex* | -.45 [-.72 -.21] | 3.47 |
| *Superior Frontal Gyrus* | -.43 [-.67 -.18] | 3.39 |
| *Precuneus* | -.33 [-.57 -.10] | 2.79 |
| *Putamen* | -.33 [-.57 -.09] | 2.73 |
| *Hippocampus* | -.29 [-.50 -.08] | 2.67 |
| *Insula* | -.31 [-.53 -.08] | 2.64 |
| *Frontal Pole* | .52 [.28 .76] | 4.30 |
| *Pericalcarine Cortex* | .46 [.24 .68] | 4.06 |
| *Heschl's Gyrus* | .42 [.18 .66] | 3.44 |
| *Parahippocampal Gyrus* | .39 [.15 .63] | 3.24 |
| *Caudal Part of Middle Frontal Gyrus* | .34 [.09 .59] | 2.69 |
| *Superior Temporal Gyrus* | .36 [.10 .63] | 2.67 |
| *Lingual Gyrus* | .34 [.09 .59] | 2.66 |
| *Supramarginal Gyrus* | .33 [.07 .60] | 2.46 |

In order to describe succinctly the following results, we define two types of GMV alterations at a lower hierarchical level. Those that reflect the direction of alterations at a higher level will be referred to as *concordant* GMV alterations, for instance larger GMV at a lower level relative to larger GMV at a higher level. Conversely, *discordant* alterations will refer to cases when there was larger GMV at a lower level relative to smaller GMV at a higher level or smaller GMV at a lower level relative to larger GMV at a higher level. Within each class of alterations, a further subdivision can be made: those associated with significant between-group difference in raw GMV (i.e., *absolute* larger/smaller GMV) and those showing no significant between-group difference in raw GMV (i.e., larger/smaller GMV *relative* to a larger/smaller GMV at the higher hierarchical level). To distinguish between *relative* and *absolute* alterations in GMV, we performed complementary post-hoc analyses on raw regional and modular GMV.

Figure 3*A* shows the hierarchical relations between GMV alterations. Very preterm born individuals had an overall smaller global GMV compared to controls, but this pattern was not uniformly distributed across different modules and brain regions. At the modular level, further smaller concordant GMV was observed in very preterm individuals in subcortical/medial temporal module, whereas larger discordant relative GMV was observed in fronto-striatal and lateral parieto-temporal modules.

**Figure 3.**
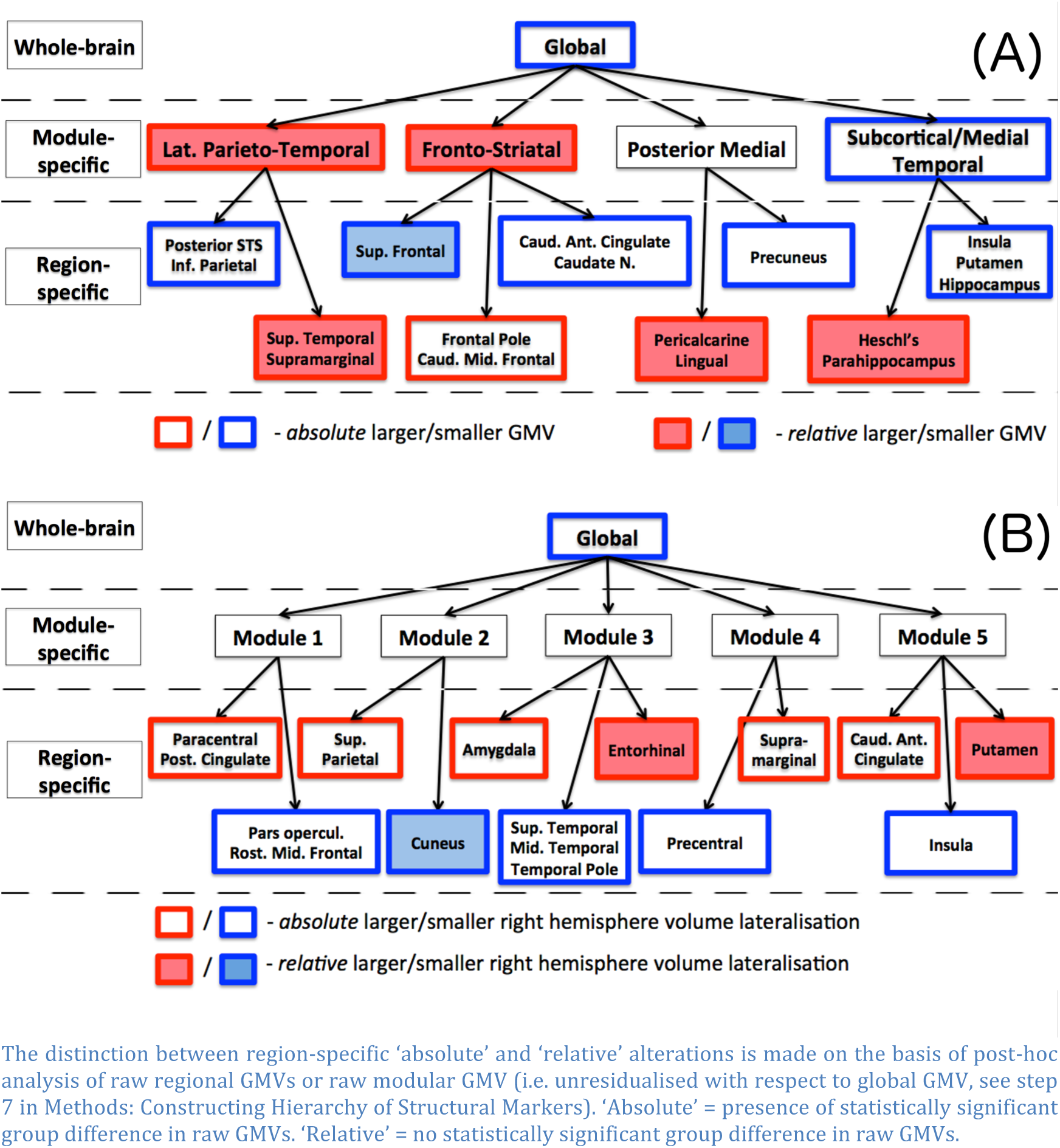
Hierarchical relations among structural alterations in very preterm individuals compared to controls. (a). Grey matter volume (GMV). (b). Grey matter hemispheric lateralisation (LI).

A range of GMV region-specific alterations was also observed in preterm individuals compared to controls. These alterations can be summarised into four groups:

1) **Smaller GMVs, concordant with alterations at a higher hierarchical level (either global, modular or both).** Absolute alterations of this type were observed in precuneus (medial posterior module), and three regions in the subcortical/medial temporal module: insula, putamen and hippocampus.
2) **Larger GMVs, discordant with alterations at a higher hierarchical level (both global and modular).** Relative alterations of this type were observed in Heschl’s and parahippocampal gyri (subcortical/medial temporal module) and pericalcarine and lingual cortices (medial posterior module).
3) **Smaller GMV, concordant with alterations at the global level, but discordant with respect to larger GMVs at the modular level.** Absolute alterations of this type were observed in inferior parietal cortex (parietal inferior lobule without supramarginal gyrus(Desikan et al., 2006)) and posterior part of superior temporal sulcus (both lateral parieto-temporal module), anterior cingulate gyrus and caudate nucleus (all fronto-striatal module). Smaller relative GMV was found in the superior frontal gyrus.
4) **Larger GMV, concordant with alterations at the modular level, but discordant with respect to smaller global GMV**. Absolute alterations of this type were found in frontal pole and caudal part of middle frontal gyrus (both in fronto-striatal module). Larger relative GMV was observed in superior temporal and supramarginal gyri (both in lateral parieto-temporal module).

Three regions also showed a significant interaction between group and time of assessment (Figure 4*A*). Very preterm individuals showed smaller GMV compared to controls by adulthood in lingual gyrus and amygdala. On the contrary, GMV in the caudal part of anterior cingulate gyrus was was smaller in very preterm individuals compared to controls at first two assessments (adolescence and early adulthood) only.

**Figure 4.**
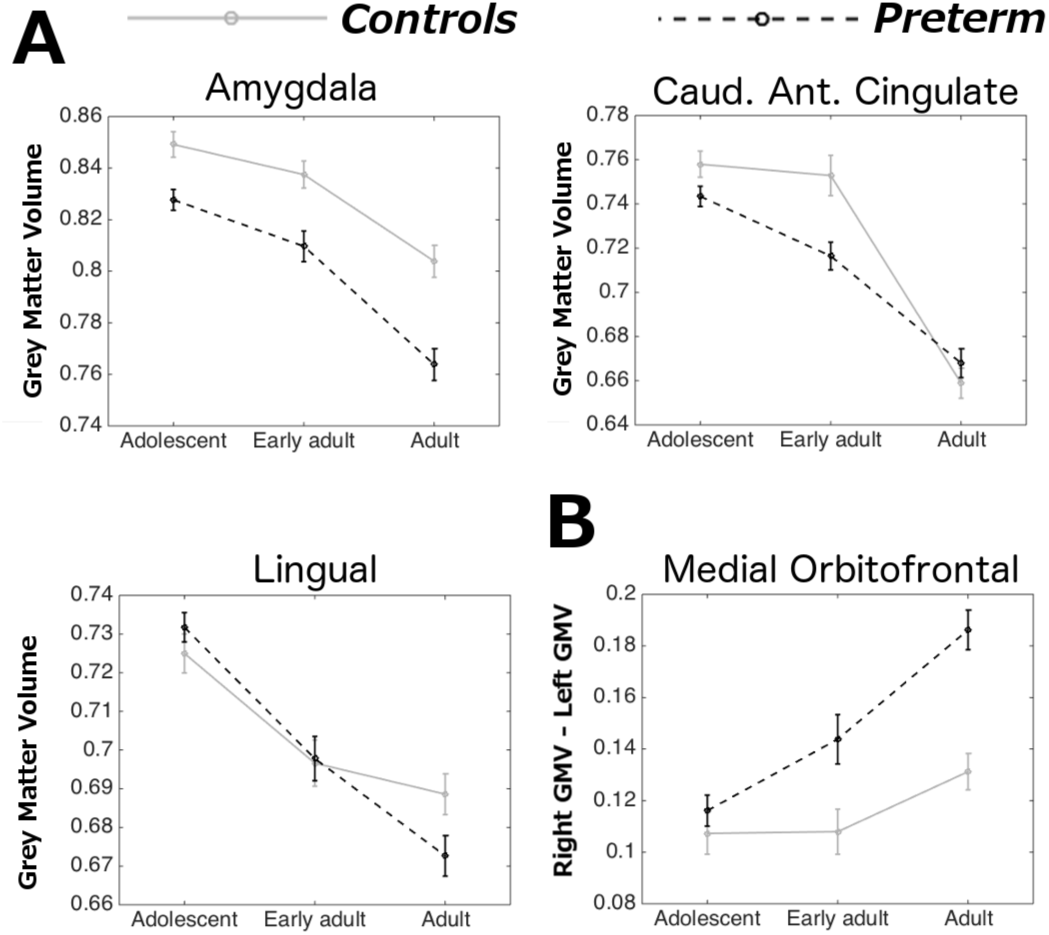
Time by group interactions. (a) Raw grey matter volume (GMV) (b) Raw lateralisation index (LI). Positive values indicate larger volume in the right hemisphere. Bars indicate standard error of the mean.

### Alterations in grey matter hemispheric lateralisation

Results of mixed effect model fitting the main effect of group on LI are shown in Table 3. Significant between-group differences in LI were observed at the global level, with preterm born individuals showing smaller right lateralization than controls. A post-hoc analysis on raw GMV showed that these results were driven by an overall smaller right GMV in the preterm group (t = 3.26, p < .005). No module-specific LI alterations were observed in the preterm group compared to controls, although there were a number of region-specific LI alterations. The hierarchical structure of these alterations is shown in Figure 3*B*. With respect to alterations observed at the global level, these can be characterised as:

**Table 3.** Summary of LI alterations in very preterm individuals compared to controls.

|  | **Standardised $\beta$ [95% CI]** | **t (df = 552)** |
| --- | --- | --- |
| **Global** | -.61 [-.86 -.36] | 4.82 |
| **Region-specific** |  |  |
| *Superior Temporal Gyrus* | -.43 [-.68 -.18] | 3.44 |
| *Pars Opercularis* | -.42 [-.67 -.18] | 3.41 |
| *Cuneus* | -.40 [-.64 -.16] | 3.33 |
| *Middle Temporal Gyrus* | -.38 [-.62 -.14] | 3.07 |
| *Temporal Pole* | -.39 [-.65 -.14] | 3.00 |
| *Rostral Part of Middle Frontal Gyrus* | -.37 [-.63 -.11] | 2.81 |
| *Insula* | -.30 [-.52 -.09] | 2.76 |
| *Precentral Gyrus* | -.31 [-.54 -.09] | 2.72 |
| *Caudal Part of Anterior Cingulate Gyrus* | .69 [.44 .93] | 5.46 |
| *Amygdala* | .52 [.30 .75] | 4.59 |
| *Supramarginal Gyrus* | .57 [.32 .82] | 4.51 |
| *Paracentral Gyrus* | .47 [.22 .72] | 3.70 |
| *Posterior Cingulate Gyrus* | .43 [.18 .68] | 3.37 |
| *Entorhinal Gyrus* | .43 [.18 .69] | 3.31 |
| *Superior Parietal Gyrus* | .40 [.16 .65] | 3.28 |
| *Putamen* | .31 [.08 .55] | 2.60 |

**Negative beta-values indicate a relative volume decrease in the right hemisphere in the very preterm group.**

1) **Concordant smaller right lateralisation** (i.e., a decrease in right GVM relative to left GMVs in the very preterm group compared to controls). Absolute alterations of this type were found in the pars opercularis, middle frontal gyrus, superior and middle temporal gyri, precentral gyrus and insula. Relative alterations of this type were found in cuneus.
2) **Discordant larger right lateralisation.** Absolute alterations of this type were observed in paracentral, posterior cingulate, superior parietal, amygdala, supramarginal cortices and caudal part of anterior cingulate gyrus. Relative alterations of this type were found in enthorinal cortex and putamen.

One region only, the medial orbitofrontal cortex, also showed a significant interaction between group and time of assessment (**Figure 4*B***). For this region, both groups showed larger right GMV, but in the very preterm group the difference between right and left medial orbitofrontal GMV increased with age significantly faster than in controls.

### Accounting for SES and handedness

When mixed-effect models for GMVs and LIs were re-run including SES at adulthood as an additional predictor, all results remained unaltered. The results for LI accounting for the effect of handedness are presented in the **Supplementary Materials.**

### Alterations in module-specific GMV and full-scale IQ

As a proof of concept that discordant relative increases may implicate compensatory mechanisms, we tested whether alterations in module-specific GMVs showed a stronger association with full-scale IQ in the very preterm group compared to controls (see Methods for details of modelling). There was a significant interaction between group and lateral parieto-temporal GVM (β = 1.27 points/standard deviation, CI = [.47 - 2.06], t(441) = 3.17, p = .0075, Bonferroni-corrected), showing that larger GMV in this module was more strongly associated with higher IQ in the very preterm group compared to controls.

### Maturation analysis

The correlation between age predicted by LASSO regression model and chronological age in the normative data set for GMVs only, LIs only and GMVs and LIs combined were, respectively, .81, .42 and .81 (Figure S1, **Supplementary Materials**). These values were comparable to the correlations obtained by support vector regression using raw GVMs and LI as features (.82, .36, and 80, respectively). This result indicates that the GMVs-only feature set represents an optimal set for estimating participants’ predicted age.

Mixed-effect analysis of maturation indices (MI) revealed significantly higher MI values in very preterm participants compared to controls (β = 4.04, CI =[2.55 5.54], t(550) = 5.31, p < .001), with no significant interaction between time of assessment and subject group (F(2, 550) = 1.15, p = .32).

The contributions of each LASSO-selected feature to group differences in MI are shown in Figure 5. Alterations at the global level, in the parieto-temporal, subcortical/medial temporal and fronto-striatal modules, as well as alterations in the frontal pole, contributed to a higher MI in the very preterm group. On the other hand, region-specific alterations in the primary auditory and visual cortices, inferior parietal gyrus and caudate nucleus were associated with delayed maturation in preterm individuals.

**Figure 5.**
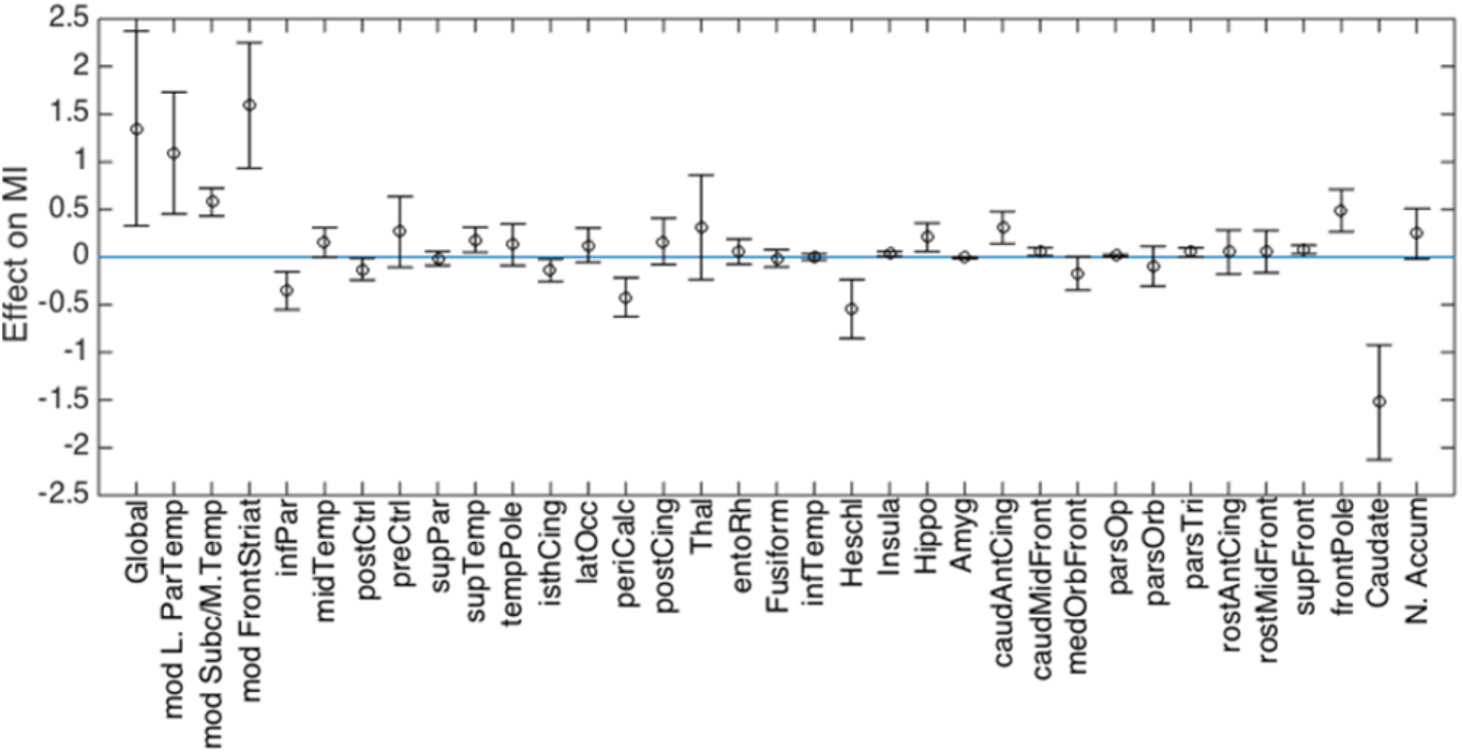
Contribution of GMV features to differences in MI between very preterm individuals and controls. Positive values indicate a GMV values contributing to a MI increase in the very preterm group compared to controls (i.e. older age); negative values indicate a GMV values contributing to a MI decrease in the very preterm group compared to controls (i.e. younger age).

We also investigated whether MI was associated with neonatal risk factors associated with very preterm birth: gestation age (GA) and ultrasound classification. The effect of ultrasound classification on MI was not significant (p = .20), but there was a significant negative association between MI and GA (t(316) = − 2.34, p = .02). Given that predicted participants’ age represents a linear combination of GMVs weighted by coefficients of the age-predictive model, we also confirmed that its association with GA was very specific and could not be accounted for by collinearity between MI and some arbitrary linear combination of GMVs (see **Supplementary Materials** and Figure S2).

## Discussion

In the present study we developed a novel analytic approach to study volumetric grey matter alterations in adults who were born very preterm as part of a hierarchically stratified system, considering patterns of anatomical co-variance. Firstly, we showed that widespread volumetric grey matter differences between very preterm individuals and controls that were observed in adolescence (Nosarti et al., 2008; Nosarti et al., 2014) mostly persisted into adult life. Such differences included a global GMV reduction in the very preterm group, consistent with prior findings (Bjuland, Rimol, Lohaugen, & Skranes, 2014; de Kieviet et al., 2012; Nosarti et al., 2002). However, GMV reduction was not uniform, and was especially pronounced in subcortical and medial temporal regions, accompanied by a relative GMV increase in frontal and lateral parieto-temporal cortices. Furthermore, global GMV reduction was greater in the right compared to the left hemisphere. Such findings suggest that perturbation of typical brain development, in this case following very preterm birth, can result in a cascade of life-long brain alterations.

Secondly, we related spatial GMV alterations to the process of brain maturation (Cole, Leech, Sharp, & Alzheimer’s Disease Neuroimaging, 2015; Franke et al., 2012) and found that adolescents and adults born very preterm had “older-appearing” brains than controls. We also demonstrated a specific association between accelerated maturation in the very preterm sample and lower gestation age.

### Regional alterations in very preterm individuals and their functional implications

We identified GMV alterations which were either in line with (referred to as ‘concordant’) or contrary to (referred to as ‘discordant’) alterations seen at a higher hierarchical level. This allowed us to demarcate which anatomical structures were more vulnerable or more resilient to the long-term consequences of very preterm birth, possibly reflecting neuroplastic adaptation. In some cases, detailed below, discordant alterations may reflect compensatory mechanisms.

Discordant GMV increases in fronto-striatal and lateral parieto-temporal modules suggest that these regions show resilience to the impact of preterm birth, as seen in global GMV reduction. There may be two possible explanations for this. First, these higher cognitive areas are involved in integrating affective, motor, and sensory responses in the appraisal and generation of complex behavioural outputs (Mesulam, 1998), and may be more affected by environmental than by perinatal factors. Second, these regions lie far from the lateral ventricles, in contrast to the subcortical and medial temporal cortices, which are vulnerable to perinatal brain injuries related to preterm birth (Volpe, 2009). In line with this, our analysis demonstrated module-specific (additive) GMV reduction in the subcortical/medial temporal module, concordant with patterns observed globally. We previously noted a lateral-to-ventral gradient of structural vulnerability while investigating the impact of the perinatal brain injury on adult structural connectivity in a subsample of the current participants (Froudist-Walsh et al., 2015; Karolis et al., 2016).

Within a module, regions of higher vulnerability and resilience could also be identified. The regions displaying the greatest vulnerability in the fronto-striatal and lateral parieto-temporal modules were likely to be those which showed a discordant GMV reduction relative to (possibly, compensatory) modular increases and included the inferior parietal cortex, anterior cingulate gyrus and caudate nucleus. On the other hand, superior temporal and supramarginal cortices, by showing a relative GMV increase, appeared as being resilient to global GMV reduction. Alterations in two regions that are part of the fronto-striatal module, namely frontal pole and caudal part of middle frontal gyrus, were somewhat difficult to characterise as they showed a GMV increase in absolute terms. With the current analyses we could not ascertain whether these results reflected a different type of vulnerability (e.g., delayed cortical thinning) or region-specific over-compensation for GMV reduction. Within the subcortical/medial temporal module, further region-specific GMV reductions were found in putamen, hippocampus, and insula, which indicate sites of preferential vulnerability to preterm-related brain injury. Within the medial posterior module, which did not show module-specific alterations, region-specific volume reduction was found in precuneus.

Unbalanced hemispheric development characterises certain developmental disorders (Zhao, Thiebaut de Schotten, Altarelli, Dubois, & Ramus, 2016) and could underlie alterations in functional connectivity lateralisation of the very preterm brain (Kwon et al., 2015; Scheinost, Lacadie, et al.,2015). However, LI alterations observed in the present study were difficult to summarise quantitatively. There was only limited evidence, after accounting for handedness (see **Supplementary Materials**), that the module comprising the basal ganglia and anterior cingulate gyrus in very preterm individuals was resilient to the pattern observed globally, which was smaller right lateralization compared to controls. Meanwhile, region-specific LI alterations in the very preterm group demonstrated a rich spatial diversity. Region-specific increase in left lateralisation, concordant with global LI alterations, was predominantly observed in lateral temporal and frontal regions, and the insula. A discordant increase in right lateralisation was observed in medial and parietal regions, and amygdala.

Specific hypotheses on the functional significance of the observed GMV alterations could be proposed for future research. Several of the regional variations in grey matter development observed in very preterm individuals seem to overlap with networks defined using resting-state functional connectivity. For example, they showed regional GMV reductions in regions that form the ‘salience’ (striatum, anterior cingulate, insula)(Seeley et al., 2007), and the ‘default mode’/‘episodic memory’ network (precuneus, medial prefrontal cortex, posterior cingulate gyrus, inferior parietal cortex and temporal regions)(Raichle et al., 2001; Spreng & Grady, 2010). These results suggest a possible link between volumetric grey matter developmental trajectories and functional connectivity. That these regions in particular show the greatest regional GMV reductions in the very preterm brain is striking, as we previously reported that the very preterm and control groups could be accurately classified on the basis of functional connections between these networks, with the salience-default mode cortico-striatal connections contributing most to this classification (White et al., 2014). Several regions from these networks have also been implicated in emotion regulation and psychiatric disorders (Shepherd, Laurens, Matheson, Carr, & Green, 2012; Townsend & Altshuler, 2012), and their altered development may underlie a higher prevalence of psychopathology in very preterm samples (Burnett et al., 2011; Nosarti et al., 2012), as well as of emotion regulation problems (Woodward, Lu, Morris, & Healey, 2017).

On the other hand, very preterm individuals may rely to a greater extent on their ‘resilient’ brain structures to perform specific cognitive operations. As a proof of concept we explored the association between modular GMV and full-scale IQ and demonstrated a stronger association between larger lateral parieto-temporal GMV and full-scale IQ in very preterm individuals than in controls. Recently, we have also demonstrated that frontal executive functions (Burgess & Shallice, 1996) played a disproportionally greater role for well-being and real-life achievement in the current very preterm-born sample compared to controls (Kroll et al., accepted).

Finally, several regions showing either region-specific GMV increases or alterations in hemispheric lateralisation have been implicated in different aspects of language processing. GMV increases include the superior temporal, supramarginal, lingual, and Heschl’s gyri, whereas a greater left hemisphere lateralisation (or smaller right lateralization) is seen in the pars opercularis, superior temporal and middle temporal gyri. Even though one can only speculate on the functional significance of these alterations, prior studies suggested that compensatory mechanisms may support cognitive and language processing in very preterm samples (Luu, Vohr, Allan, Schneider, & Ment, 2011; Scheinost, Lacadie, et al., 2015).

### How structural alterations reflect maturational processes

Several researchers proposed the hypothesis of delayed brain maturation at earlier stages of development in survivors of very preterm birth (Back & Miller, 2014; Back et al., 2007; Pierson et al., 2007). The present study found little evidence to support this hypothesis when grey matter development from adolescence and beyond is concerned. Contrary to expectations, the observed structural characteristics of the very preterm brain suggested accelerated rather than delayed maturation. However, this pattern was not uniformly distributed across different hierarchical levels. Whereas global and modular structural markers contributed positively to a higher maturation index, a few regions, most notably the caudate nucleus, demonstrated patterns of delayed maturation. Importantly, accelerated maturation was associated with younger gestational age but not with severity of perinatal brain injury, in line with suggestions that subtle (diffuse) forms of developmental alterations (Volpe, 2009), which do not entail acute neuronal loss, determine the course of neurodevelopment following very preterm birth (Back & Miller, 2014).

In this study we also failed to find convincing evidence for a developmental ‘catch-up’ we previously suggested (Nam et al., 2015). Only a few region-specific GMVs and LIs showed significant between-group differences in longitudinal changes. In 3 out of 4 cases between-group differences increased with time. We cannot rule out the possibility that ‘catch-up’ might have occurred in a limited number of regions by the time of adolescent assessment. For example, we did not find strong evidence that thalamic volume was disproportionally affected in the studied sample, whereas reduction in thalamic volume and connectivity has been a consistent finding in very preterm infants (Ball et al., 2013; Ball et al., 2012). However, the indirect effect of early alterations in thalamic development may still persist. Five out of 7 cortical regions that were identified in the present study as showing independent region-specific GMV increases in the very preterm brain, namely superior temporal, supramarginal, lingual, Heschl’s and pericalcarine gyri, were also reported as having reduced thalamocortical connectivity in infants (Ball et al., 2013).

A revision of the concept of brain “maturation” is perhaps required to reconcile the two lines of evidence, i.e., delayed maturation at earlier stages and accelerated ageing from the time of adolescence onwards. A formula ‘less volume – older brain’, despite being useful in numerous contexts, is too simplistic to characterise age-dependent developmental trajectories as it ignores their non-linear patterns, the diversity of their functional forms (Douaud et al., 2014) and their mutual interactions. In this context, ‘dysmaturation’ or ‘under-development’ (Back & Miller, 2014), which we define as an inability to reach predetermined peaks in typical developmental trajectories, may be more appropriate terms to use. Unlike ‘delayed maturation’, the two terms do not imply that the very preterm brain is always lagging behind a control brain. Given the non-linearity of GVM development, a dysmaturation at earlier stages of development may lead to an accelerated ageing at a later stage. For instance, in healthy controls, existing evidence suggests that a grey matter pre-pubertal increase in a number of cortical regions is followed by a post-pubertal decrease, with changes accompanying maturing cognitive abilities (Giedd et al., 1999; Gogtay et al., 2004). Consequently, smaller global GMV in very preterm individuals during the pre-pubertal period can be interpreted as a maturational delay. However, post-puberty onwards, a decreased grey matter volume can be interpreted as an acceleration of typical brain maturational processes, i.e., accelerated ageing.

The functional consequences of unbalanced maturation require further investigation. Of relevance, a prominent model of neurodevelopment proposes a relative faster maturation of the striatum during adolescence compared to the slower developing prefrontal cortex, which is associated riskier behaviours (Somerville & Casey, 2010). In contrast, adolescents born very preterm tend to display less risk-taking behaviour than term-born controls (Saigal, 2014). Our results suggest a potential biological mechanism to explain the relatively reduced risk-taking behaviour. In individuals born very preterm, in contrast to controls, prefrontal cortical structures show greater signs of maturity while the caudate nucleus of the striatum is relatively immature. This may signify that a pattern of “asynchronous development” of these structures that is seen during typical development (Somerville & Casey, 2010) does not occur following very preterm birth.

### Study limitations

One of the obvious study limitations, endemic to neuroimaging long-term longitudinal data, is the varying scanner and scanner parameters between different times of assessment. Even though the possibility of this factor affecting the results cannot be completely excluded, we believe it may have had only a limited impact on the present results for the following reasons: first, at each cross-sectional assessment, scanning parameters were identical in very preterm individuals and controls; second, the effect of scanner across all groups was modelled explicitly in our analyses by introducing time of assessment as a categorical variable of no interest. Finally, the interaction between group and time would be most susceptible to these effects, whereas most of our findings showed a main effect of group only. A variable pattern of significant interactions between different regions makes it unlikely that these resulted as a consequence of a systematic bias due to scanning parameters.

## Conclusions

We characterised structural alterations in the very preterm brain and then used such alterations as markers of age-dependent maturational processes. We studied grey matter volumes as part of a hierarchically stratified system and showed widespread and stable (from adolescence to adulthood) volumetric grey matter differences between the very preterm and the control groups. These highlighted brain areas that are both vulnerable and resilient to the long-term consequences of very preterm birth.

The finding of increased maturation indexes in the very preterm brain emphasizes the importance of future studies on the ageing process in very preterm samples. It remains to be investigated whether these quantitative differences have real-life implications, for instance whether early brain ageing is associated with cognitive decline, as observed in traumatic brain injury (Cole et al., 2015). Such studies could inform the development of cognitive and behavioural interventions aimed at boosting brain resilience.

## Materials and Methods

### Local site participants

Two hundred and nine very preterm participants were recruited from a sample of individuals born before 33 weeks of gestation between 1979 and 1984 and admitted to the Neonatal Unit of University College London Hospital (UCLH). 188 full-term controls matched for year of birth were also studied. Controls were recruited either at birth as they were delivered at term at UCLH (n = 62), or from community advertisements in the press (n= 126). For all participants, exclusion criteria were any history of neurological complications including meningitis, head injury, and cerebral infections. Parental socio–economic status was classified according to Her Majesty’s Stationary Office Standard Occupational Classification criteria. Neonatal cranial ultrasound (US) was collected within 5 days of birth and results were summarized as (1) normal US, (2) uncomplicated periventricular haemorrhage (grade I-II), without ventricular dilatation (PVH), and (3) periventricular haemorrhage (grade III-IV) with ventricular dilatation (PVH + DIL) (Nosarti, Walshe, et al., 2011). There were no cases of periventricular leukomalacia. Assessments took place at adolescence (mean age of both groups=15.2 years), early adulthood (mean age=19.8 years) and adulthood (mean age=30.6 years). Sample statistics are shown in Table 1.

### MRI acquisition parameters

At adolescent assessment, MRI was performed at one of two sites. Data for the 1979–82 cohort and controls was acquired on a 1.5 Tesla GE Signa Horizon machine (General Electric Medical Systems, Milwaukee, WI, USA) at the Institute of Neurology, London. The 1983–84 cohort and controls were scanned using a 1.5 Tesla GE Signa N/Vi machine at the Maudsley Hospital, London. At both sites, threedimensional T1-weighted MR images were acquired in coronal plane, with a spoiled gradient recalled pulse sequence (flip angle 35°, field of view 240 mm, echo time 5 ms, repetition time 35 ms). Each image contained 124 slices with a matrix size of 256 × 256, slice thickness of 1.5 mm.

At early adulthood assessment, all participants were scanned at the Maudsley Hospital, London, with the same scanning protocol as the one used at adolescence.

At adult assessment a GE Signa HDx 3.0-T MR scanner (General Electric, USA) with an 8-channel head coil was used. T1-weighted images were acquired using an Enhanced Fast Gradient Echo 3-Dimensional (efgre3D) sequence (flip angle 20°, field of view 280 mm, echo time 2.8 ms, repetition time 7 ms), slice thickness 1.1 mm.

### Normative data set

1270 T1-weighted images of healthy control individuals, aged 9 – 65, were acquired from 5 publicly available neuroimaging databases. Details are provided in Table S1 (Supplementary Materials).

### Normalisation

A study-specific longitudinal template was created using T1-weighted images, in two steps. Firstly, three cross-sectional templates were created, one for each time point (adolescence, early adulthood and adulthood). For each template, a sample of 84 participants was randomly selected such that (a) it included, in equal numbers, subsamples of control participants, very preterm individuals with normal US and very preterm individuals with PVH or PVH-DIL; (b) any one participant could be selected for creating only one template. Images for the adolescent and early adulthood assessments were re-sliced to match the dimensions of the higher resolution images acquired at adulthood. Secondly, a study-specific longitudinal template was created using the three cross-sectional templates (Deoni, Dean, O’Muircheartaigh, Dirks, & Jerskey, 2012).

All templates were generated using the Greedy symmetric diffeomorphic normalization (GreedySyN) pipeline distributed with the Advanced Normalization Tools (http://stnava.github.io/ANTs/)(Avants et al., 2011). After an initial affine transformation, four iterations of the nonlinear template creation were performed.

Grey matter probability maps were extracted using the unified segmentation approach implemented in SPM12. These maps were non-linearly registered to the longitudinal template’s GM map using ANTS/GreedySyN. To produce GMV maps, normalised GM maps were subsequently scaled by the Jacobian determinants of the deformation field and smoothed with a Gaussian filter. As there was no need to comply with the statistical assumptions of mass-univariate analysis, a relatively small FWHM of 2 mm was used to correct for imprecisions in normalisation.

### Grey matter volume

Average GMV per voxel was calculated for 82 cerebral and subcortical regions demarcated by the automated FreeSurfer parcellation of the longitudinal template (Fischl, 2012). The measure was obtained by first creating brain regional masks, thresholded at p > .2 and then binarised. Second, these masks were multiplied by a study-specific GMV mask obtained by averaging all participants’ GMV maps that were selected for template construction, also thresholded at p > .2 and binarised. Finally, these masks were used to extract regional GMV from individual maps, which were subsequently averaged across all voxels in the mask.

### Hierarchy of structural markers

Feature re-parameterisation was performed as follows:

1) Two types of volumetric markers were obtained: (a) the mean of homologous regions in the left and right hemisphere, hereafter - regional GMVs; (b) the difference between homologous regions in the left and right hemisphere - hereafter - regional lateralisation indices (LIs). The further steps refer to GMVs, but were exactly the same for LIs.
2) GMVs were combined into two *N* x 41 feature matrices, one for the normative data set and one for the study sample, where *N* is the number of participants in a sample (either 1270 or 560) and 41 is the number of GMVs.
3) A 41x41 correlation matrix was calculated for the normative dataset, so that each element of the matrix reflected the correlation between GMVs in a pair of distinct brain regions across subjects.
4) Modularity analysis was run on the above correlation matrix, to subdivide the entire set of brain regions into modules. Modular partitioning was obtained using the Louvain method for community detection (Blondel, Guillaume, Lambiotte, & Lefebvre, 2008). As certain heuristics are embedded in the algorithm, the partitioning was performed 1000 times (Dwyer et al., 2014). A matrix was generated on each algorithm iteration, where element (*i, j*) assumed the value of 1 if the nodes *i* and *j* were designated to the same module; otherwise it assumed the value of 0. Resulting matrices were averaged across iterations to obtain a consistency matrix, which was submitted again to the Louvain process for a final partitioning into modules.
5) Following partitioning, the original feature matrices of each data set were divided into submatrices, **R_l_**, …, **R_k_**, such that any **Ri** combined the regional GMVs that belonged to a particular module, and *k* was the number of modules identified by modularity analysis.
6) A principal components analysis (PCA) was performed on the normative data set. The first principal component, **PC-a^i^**, was calculated for each module independently, i.e., for each **R_i_** in the normative dataset. We call this stage PCA-A, to distinguish it from a later stage in the analysis.
7) Individual data points (i.e., the rows of **R_i_**) in both data sets were projected onto **PC-a^i^**. Subject loadings along **PC-a^i^** were taken to represent GVM for module *i*.
8) Each Ri in both data sets was then residualised with respect to **PC-a^i^**. Each column **ri** of a residualised **Ri** was taken to represent a region-specific GMV (see Figure 1*A*).
9) − 12) Steps (5)–(9) were repeated in the next hierarchical level of analysis, PCA-B, but this time for modular GMVs (i.e., subject loadings along all PC-**a^i^’**s) which were considered as members of one, global, module. In this way, we obtained global and module-specific GMVs, **g**, and **m_i_**, respectively (see Figure 1*A*-*B*).

### Longitudinal trajectories in GMVs and LIs

Linear mixed-effect models were used to evaluate the effect of very preterm birth on longitudinal GMV trajectories. Models were fitted to each GMV and LI marker independently. All fitted models included six fixed factors: 1) intracranial volume (ICV); 2) sex; 3) group; 4) time (as a categorical variable to account for scanner differences); 5) group x time interaction; and 6) interaction between time of assessment and the deviation of a participant’s age from the mean age of participants at the time of a corresponding assessment. The latter factor, combined with a categorical time factor, would account for non-linear age effects and control for very preterm participants’ slightly older age compared to controls (see Results).

Three random effects, modelling the covariance structure of the data, were evaluated: 1) intercept only, 2) intercept and participants’ age (centred on the grand mean) combined and 3) intercept, participants’ age and participants’ age squared combined. A log-likelihood model comparison test demonstrated that the set of random factors that included intercept and participants’ age outperformed the intercept model in all but one case. It also outperformed the model that included intercept, participants’ age and participants’ age squared in 80% of cases. Consequently, the intercept and linear effect of age were retained as random factors in all subsequent statistical analyses. False discovery rate (FDR) corrections were applied to all *p*-values to account for multiple statistical testing.

### Module-specific GMVs and IQ

As a proof of concept that discordant relative increases (see Results section for an explanation of the term) may implicate compensatory mechanisms, we tested whether alterations in module-specific GMVs demonstrate a stronger association with cognitive outcome in the very preterm group compared to controls. In order to do this, we residualised module-specific GVMs with respect to the model described in the previous section. We then constructed a model that contained fixed intercept, group, sex, and group-time interaction factors, random intercept factor and (one at a time) an interaction between group and a module-specific GMV as an additional fixed factor. The model was fitted to full-scale IQ and results were Bonferroni-corrected.

### Maturation analysis

To determine whether the observed GMV alterations in the very preterm sample reflected a delayed or accelerated maturation, we used LASSO regression – a method allowing simultaneous data modelling and sparsification of the predictor set. To determine the degree of sparsification, we firstly half-split the normative data set into training and validation sets. We fitted a LASSO regression model to the training set using a range of penalty values (the greater the value – the stronger the sparsification) and then applied estimated parameters to the validation set. We determined the model likelihood for each choice of a penalty value and selected a predictor set that was associated with a maximum likelihood. We used the correlation between predicted age and chronological age as a measure of prediction accuracy. The latter was used to determine which feature set, GMVs only, LIs only, or GMVs and LIs combined, was optimal for age prediction. Next, we fitted an ordinary multiple regression model to all data in the normative sample in order to obtain more accurate estimates of beta values for the predictors that survived LASSO regularisation. These estimated model parameters were then applied to the study sample data. Participants’ chronological age was subtracted from their predicted age to obtain a maturation index (MI). Negative and positive MI values indicate delayed maturation and accelerated ageing, respectively. Because of likely biases in co-registration of normative data set to the study sample template, we treated MI estimates as having arbitrary units, i.e., without implying that a maturation index equal to 5 is equivalent to accelerated maturation by 5 years. Between-group MI difference was tested with mixed-effects linear regression using the set of predictors described in the previous section.

In the very preterm sample only, we also investigated whether MI was associated with neonatal risk factors, gestation age or severity of perinatal brain injury. The mixed-effect models used for GMVs and LI analysis were adjusted, by substituting group and group by time interaction terms with (separately) gestation age or ultrasound classification and their interactions with time.

## Acknowledgements

We thank our study participants for their continuing help. We also thank the National Institute for Health Research (NIHR) Biomedical Research Centre at South London and Maudsley NHS Foundation Trust and King’s College London for supporting the neuroimaging facilities used in our study.

## Supplementary Materials

**Table S1.**
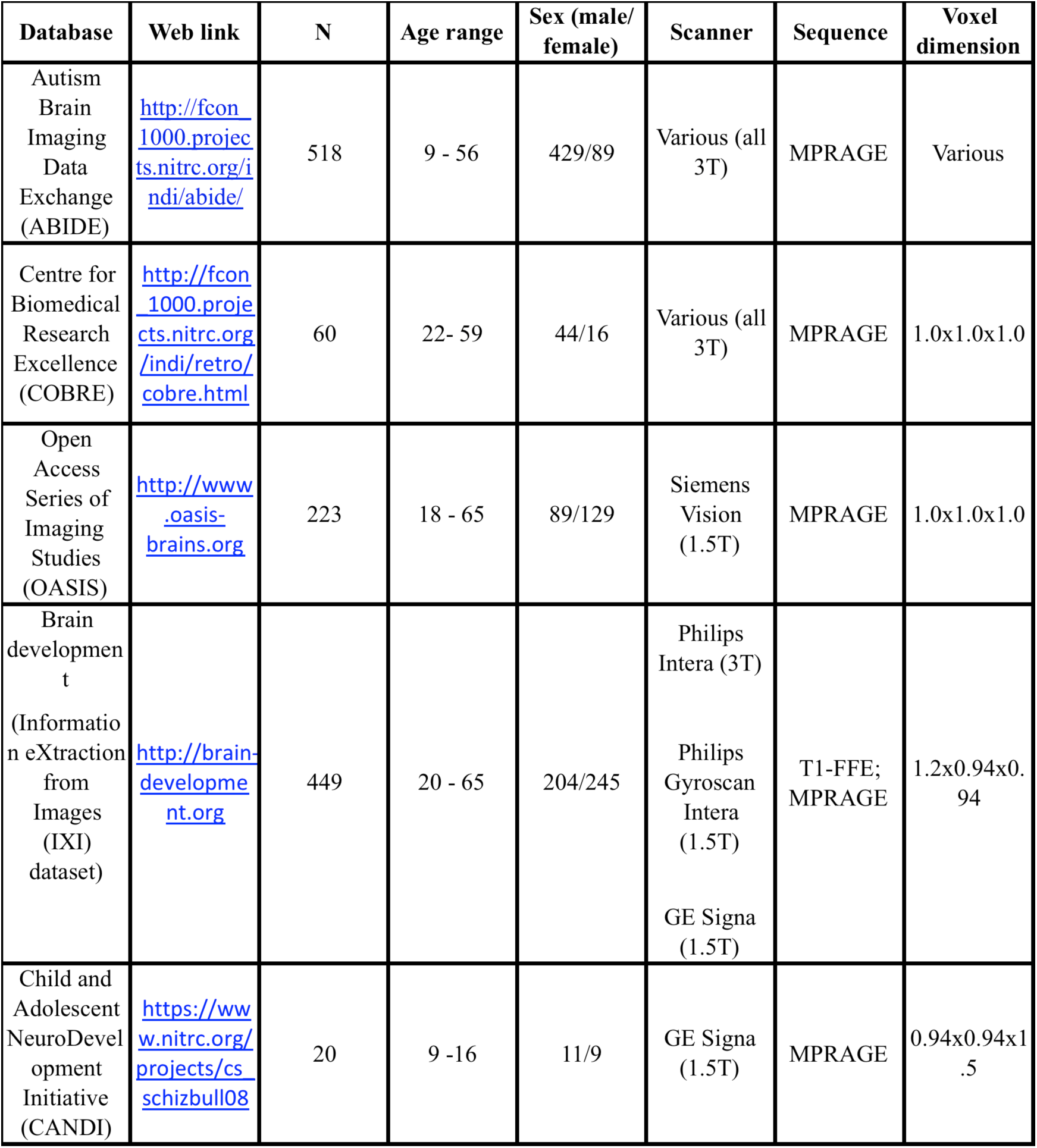
Characteristics of the normative data set

**Figure S1.**
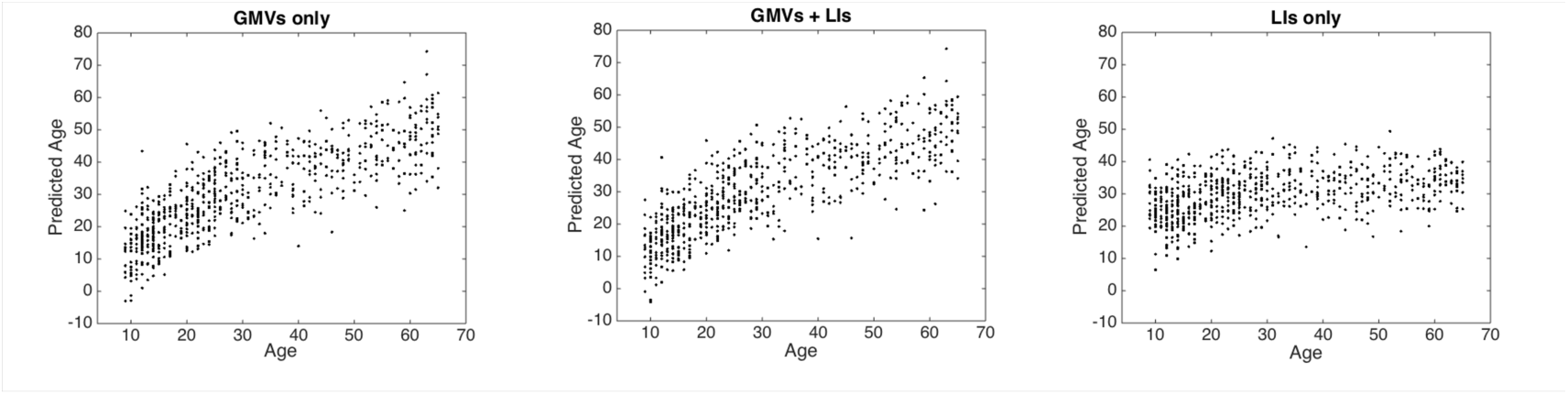
Age prediction in the normative data set (split-half) using: GMVs markers only (left panel), GMVs +LIs markers (middle panel) and LIs only (right panel).

## Handedness and LI

Even though handedness did not statistically differ between the groups, we explicitly accounted for its effect in LI analysis because of a higher prevalence of missing data in the control compared to the very preterm group. All data were included in this analysis, with missing values treated as a separate level of handedness. Results showed two discrepancies compared to the previous findings. Firstly, the relative greater right lateralisation in putamen in the very preterm group was no longer significant. Instead, Module #5, which comprised basal ganglia structures (including putamen), anterior cingulate gyrus, Heschl’s gyrus and insula, showed a relative greater right lateralisation in the very preterm group compared to controls (β = −.33, CI =[−.58 −.08]).

## Specificity of association between Maturation Index (MI) and gestation age (GA)

Given that the predicted participants’ age represents a linear combination of GMVs, weighted by coefficients of the age-predictive model, we also investigated the specificity of its association with GA, i.e. we wanted to investigate whether this association could be accounted for by collinearity between MI and some arbitrary linear combination of GMVs. To do this, we generated 20000 linear combinations of GMVs by applying random weighting in the range of −1 to 1. We then selected 1000 (5%) linear combinations, which showed the highest absolute correlation with MI. We then residualised MI with respect to each of them and fitted the models to each residualised MI in order to obtain a mean and standard deviation of regression coefficient associated with GA.

**Figure S2** shows the mean and standard deviation of standardised regression coefficients associated with GA obtained in the analysis. We also plotted the coefficients obtained after residualising with respect to a set of preselected (‘meaningful’) GMVs linear combinations. In all cases, the means of these coefficients were within 1 standard error of the coefficient fitted to the unresidualised MI.

**Figure S2.**
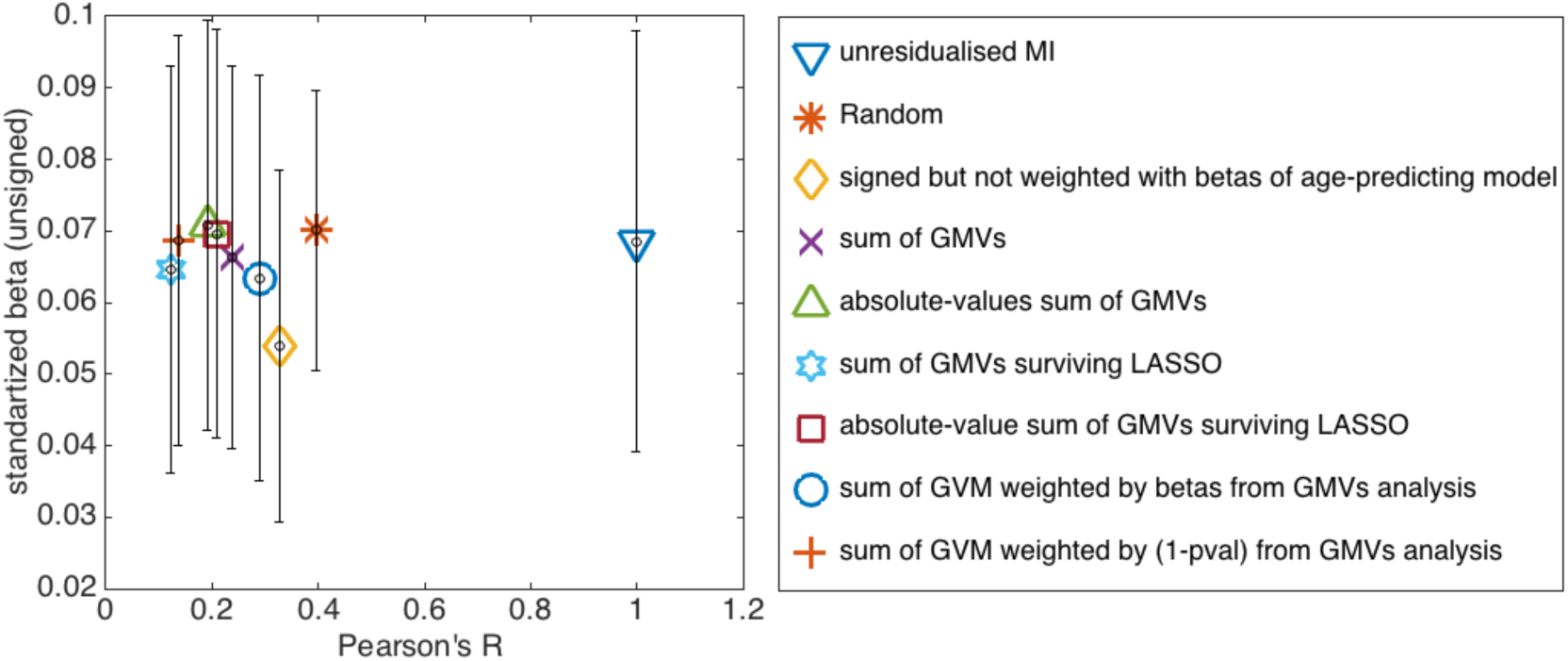
Association between GA and MI following residualisation of MI with respect to a set of GVM linear combinations, as detailed in figure legends. On x-axis, a correlation between MI and a linear combination used to residualise MI.

